# Proviral sequencing suggests the majority of the HIV reservoir is expressed over time but significant decay is obscured by clonal expansion

**DOI:** 10.1101/348409

**Authors:** Marilia Rita Pinzone, D. Jake VanBelzen, Sam Weissman, Maria Paola Bertuccio, LaMont Cannon, Wei-Ting Hwang, Brad Sherman, Una O’Doherty

**Affiliations:** Department of Pathology and Laboratory Medicine, University of Pennsylvania, Philadelphia PA 19104; Department of Biostatistics and Epidemiology, University of Pennsylvania, Philadelphia, PA 19104; Laboratory of Human Retrovirology and Immunoinformatics, Frederick National Laboratories for Cancer Research, Leidos Biomedical Research, Inc., supporting the Division of Clinical Research, NIAID; Department of Biomedical Engineering, University of Delaware

## Abstract

After initiating antiretroviral therapy (ART), a rapid decline in plasma viremia is followed by reservoir stabilization. Viral outgrowth assay suggests the reservoir continues to decline slowly, but variation over time and among individuals complicates our understanding of selective pressures during ART. We used full-length sequencing to study more than 800 HIV proviruses of two subjects on ART at four time points over nine years to investigate the selection pressures influencing the dynamics of the reservoir. We found that intact as well as defective proviruses capable of significant protein expression decrease over time. Moreover, proviruses lacking genetic elements to promote viral protein expression, yet containing strong splice donor sequences increase relative to other defectives over time, especially among clones. Our work suggests that HIV expression occurs to a significant extent during ART and results in HIV clearance, but this is obscured by clones generated by donor splice site-enhanced clonal expansion.

---

The advent of antiretroviral therapy (ART) revealed a treatment-resistant reservoir requiring life-long therapy ^1,2^. Pioneering work has shown that the HIV reservoir has a very slow rate of decay. Estimates of reservoir decay suggest a half-life of 44 months using Quantitative Viral Outgrowth Assay (QVOA) ^1–3^. However, these measurements were indirect, and their error was sufficiently large that the precise half-life of the reservoir in individual patients was uncertain. Differentiating error due to assay versus biological variation is difficult. If biological variation is prominent, a subset of subjects may have significant reservoir decline while others may have a more stable reservoir. This could be due to multiple reasons, including compliance to ART or biological differences in the host or pathogen.

Viral nucleic acid measurements have been used as a surrogate for HIV reservoir size as some studies have shown significant correlations with QVOA ^4–6^. Longitudinal studies suggest proviral DNA is relatively stable after the first few years of ART ^7^. HIV DNA measurements suffer from the presence of defective proviruses, which constitute the majority of the proviral DNA; thus, while the intra-assay variation for PCR is small ^8^ the variable and largely unknown frequency of defective proviruses results in precise but inaccurate estimates ^9–13^. As a consequence, large changes in replication-competent viruses may be masked by defective proviral DNA. Moreover, selective pressures on defective DNA may be different than selective pressures on intact proviruses ^14^ and thus HIV DNA measures may not be an appropriate way to monitor reservoir dynamics over time.

Monitoring the frequency of individual proviral sequences over time could reveal positive and negative selective pressures that might be exerted on infected cells. While such an approach is currently not feasible for all HIV-infected individuals due to limited throughput and cost, in-depth study of a subset of subjects might provide new insights into reservoir dynamics as well as the effect of the host on reservoir persistence.

We employed limiting dilution polymerase chain reaction (PCR) followed by DNA sequencing to obtain full-length sequences of integrated HIV proviruses in two subjects on suppressive ART over time. We provide evidence that defective and intact proviruses that are capable of HIV protein expression are under negative selective pressure, while defective proviruses that have weak potential for HIV protein expression, but encode a strong donor splice sequence, are under relative positive selective pressure. We also show significant biological variation in reservoir decay between and within these two individuals. We also provide evidence that clonal expansion slows decay. One optimistic implication from our analysis is that the replication-competent reservoir of intact proviruses is under more negative selection than defective proviruses and that the majority of the reservoir is expressed over time.

## Results

### Longitudinal measurement of plasma HIV RNA, total HIV DNA, integrated HIV DNA and circulating CD4 T-cell counts of two subjects started on ART (Figure 1)

We wanted to assess the decay rate of intact and defective proviruses by combining proviral sequencing with PCR measurements of HIV DNA levels. To ask this question, we identified two subjects with similar histories for whom we had peripheral blood mononuclear cells (PBMCs) aliquots over a ~8-9-year period of time after achievement of virological suppression. Total and integrated HIV DNA were assessed at 4 intervals from ~2 to 9 years after starting ART (Figure 1). The same DNA samples were utilized for proviral sequencing (Figures 2–5). Viral load and CD4 T-cell count were assessed 10 times in Subject 1 over the study period and viral load was always below the detection limit of the diagnostic assay (<50 copies/ml). Viral load and CD4 T-cell count were assessed 25 times in Subject 2 over the same time interval. Viral load was below the detection limit for all but two time points between the first and second sample, where the plasma viral load was 51 and 99 copies/ml, but was well controlled thereafter. There was a slight decline in total and integrated HIV DNA over the study period, but less than a 2-fold change by any measure (i.e. normalized to CD4, PBMC or per unit volume). In conclusion, RNA and DNA measurements including viral load, total and integrated HIV DNA decreased minimally over 7 years.

**Figure 1.**
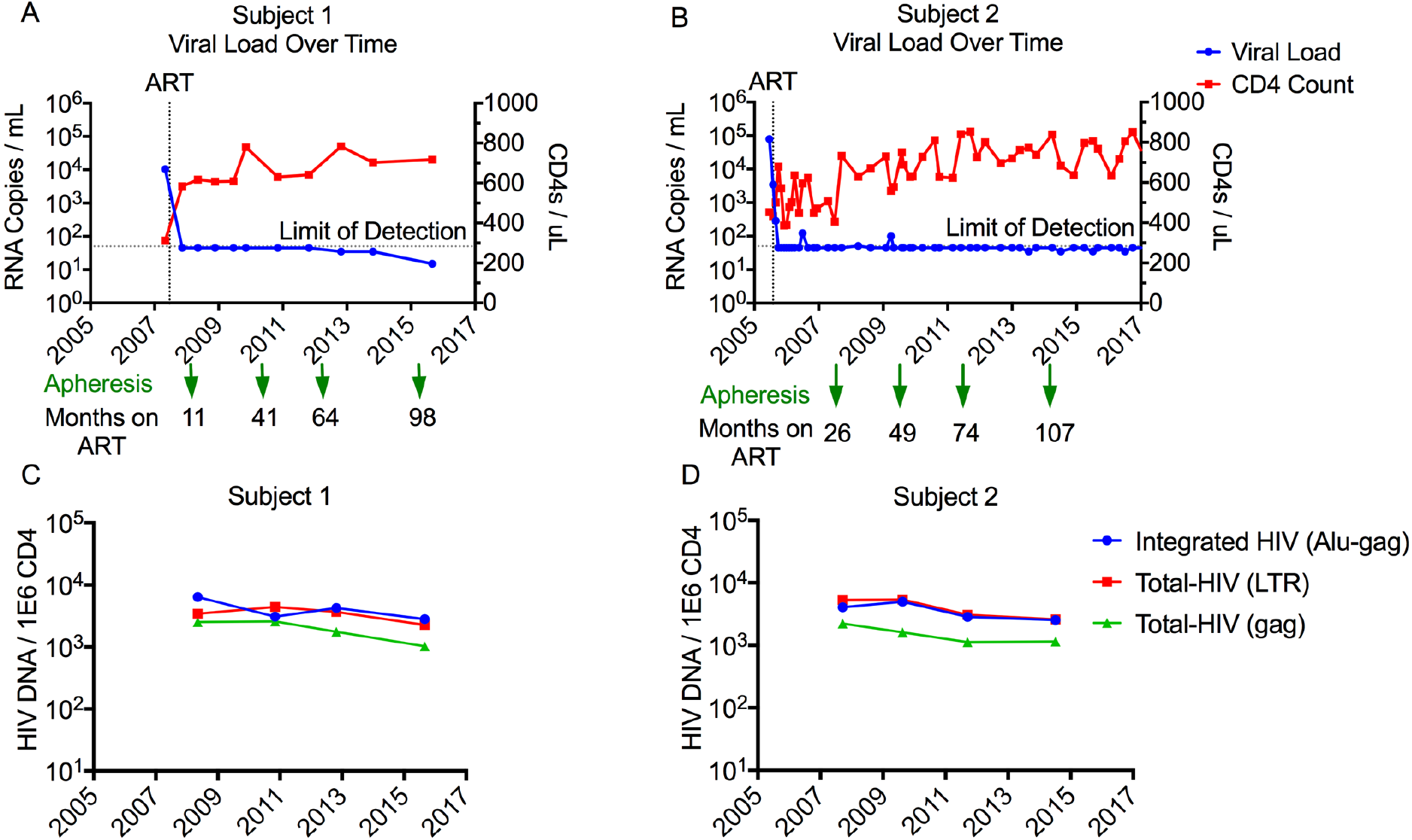
Longitudinal measurement of plasma HIV RNA, total HIV DNA, integrated HIV DNA and circulating CD4 T-cell counts of two Subjects started on ART. A-B) Longitudinal levels of plasma HIV-1 RNA and CD4 T-cell counts. For each subject, PBMCs were collected by apheresis at the four time points indicated in the graph. C-D) HIV DNA quantification by qPCR of PBMC DNA from the indicated apheresis collections. Total HIV DNA was quantified by primers bounding to the LTR region (red) or gag region (green) of HIV-1. Integrated HIV was measured by Alu-gag PCR (blue). Values are normalized to CD4 T-cell count and presented as log copies of HIV per million cells.

**Figure 2.**
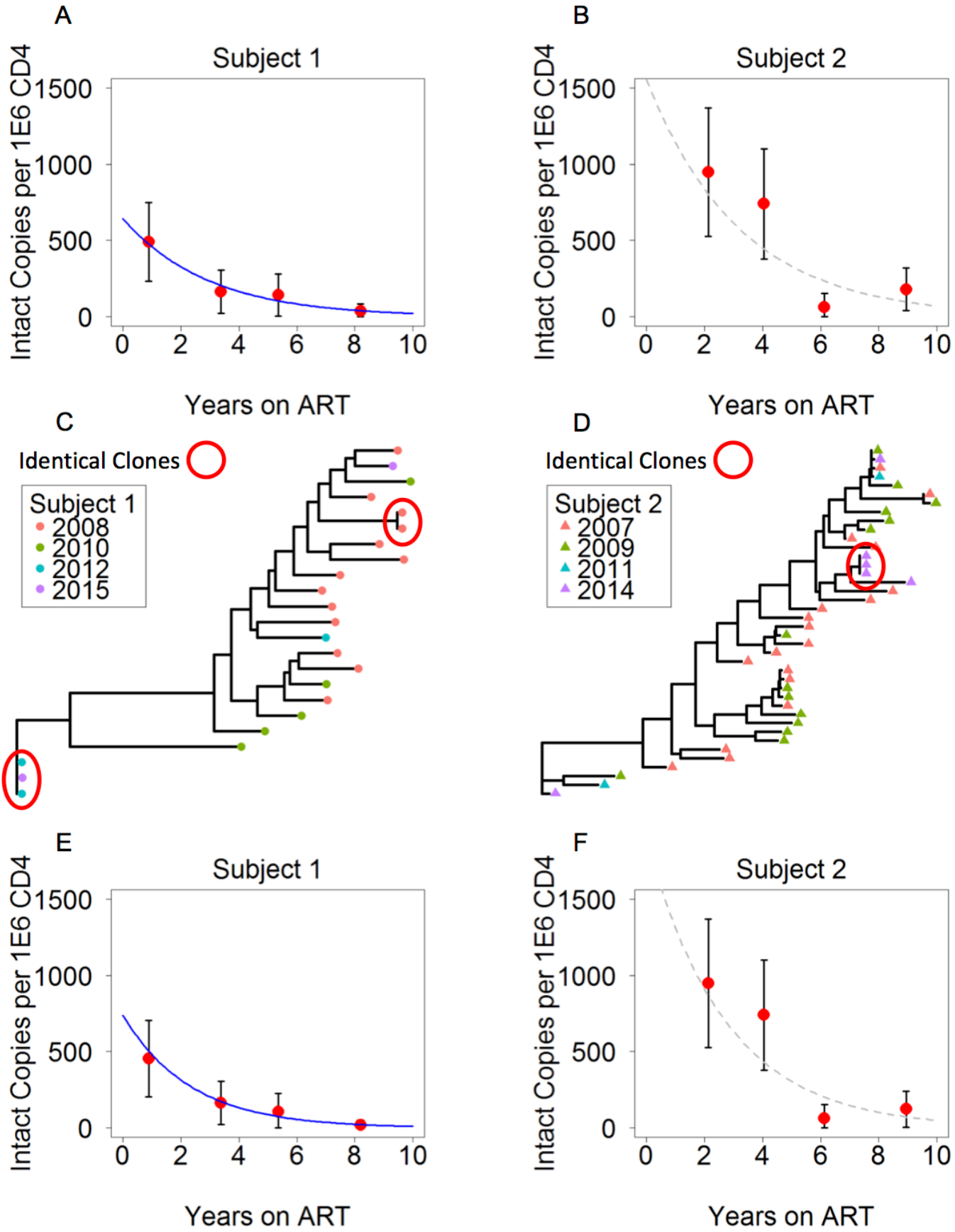
Dynamic changes of proviral characteristics over time. A) Frequency of intact proviruses throughout treatment for Subject 1 measured by intact copies per million CD4 T cells. Red marker represents calculated intact proviruses based on the concentration of sequenced proviruses that were intact and total CD4 T cell count. Black bars signify 95 percent confidence interval based on a binomial process. The blue line is the estimated decay based on a mixed effects regression model. B) Frequency of intact proviruses in Subject 2. Grey dotted line is estimated decay based on our model, which did not prove to be statistically significant. C) Phylogenetic tree of intact proviruses for Subject 1. Branch lengths are proportional to genetic distance to a consensus sequence for the sequences graphed in the tree. Identical clones are indicated within the red circles. Consensus sequences were generated for each Subject and used to root the tree. D) Phylogenetic tree of intact proviruses for Subject 2. Circled clones represent 50 percent of intact proviruses from the last apheresis collection. E) Frequency of intact proviruses for Subject 1. In order to minimize the effects of clonal expansion, clones were only recognized the first time in which they were detected. F) Frequency of intact proviruses in Subject 2 with clones counted only once, when they first appeared.

### Dynamic changes of proviral characteristics over time (Figure 2)

We performed full-length proviral sequencing at limiting dilution from 4 blood samples collected by apheresis from each subject over an 8-9 years period since starting ART. We sequenced over 800 individual proviruses and obtained approximately 100 proviruses per time point (Supplemental Figure 1). Mixed reactions containing ≥ 2 proviruses were excluded from analysis as detailed in our methods. We performed *de novo* assembly to generate contiguous sequences. Evaluation of the *de novo* assembled proviruses showed that the proviral reservoir largely consists of virus harboring deletions, as determined by size, in agreement with other studies ^9–14^.

### Dynamic changes of intact proviruses suggest stronger selective pressure against the replication-competent reservoir relative to defective proviruses (Figure 2)

To estimate the decay rate of the HIV reservoir, we counted the proviruses that appeared to be fully intact. Our criteria for an intact provirus were the presence of 9 open reading frames (ORFs), 3-4 stem loops at the psi packaging site, as well as several critical donor and acceptor splice sequences ^15–17^ and Rev-responsive element (RRE) sequence ^18–20^ as detailed in the methods section. When we plotted the frequency of intact proviruses over time, we noticed substantial decay (Figure 2A), in contrast to total or integrated HIV DNA, which were relatively constant over the same time frame (Figure 1C) (14-fold drop in intact proviruses vs 1.5-fold in total HIV DNA for Subject 1; 5.3-fold vs 2-fold drop in Subject 2). We performed an ordinary least squares fit of our data using a linear model. We then fit the data to an exponential model, and by Akaike’s Information Criterion (AIC) we found that our data was best characterized by the exponential model (AIC = 4.36, Subject 1) as compared to the linear model (AIC = 51.30) for Subject 1.

For Subject 1, we found the exponential decay rate was -0.339/year with a half-life of 2 years. For the purpose of modeling, time 0 was the moment the subject was placed on ART. Using the best fit exponential decay curve, we predicted that the subject had a concentration of approximately 640 intact proviruses per million CD4 T cells at the time he was placed on ART, which dropped to 36 intact proviruses at the last time point on ART. Subject 2 had a less straightforward decay curve, as he had an increase in the frequency of intact proviruses from 64 to 180 copies per million CD4 T cells at the fourth time point. Given the increase in intact proviruses at the last time point, exponential decay is not a good model for this Subject. It can only be said that Subject 2 has a slower overall decay. This slower overall decay of intact proviruses may be due clonal expansion, ongoing replication, redistribution of infected cells from lymphoid tissue or sampling error.

### Phylogenies of Intact proviruses suggest a role for clonal expansion in HIV persistence (Figure 2C-F)

To investigate the role of clonal expansion in the proviral decay over time, we aligned intact proviruses and generated a phylogenetic tree for both subjects independently (Figure 2C and D). We noticed that there was no increase in sequence diversity over time, consistent with no replication ^21–25^ or minimal replication ^26–31^ on ART. We identified a few identical sequences in the intact tree. For Subject 1, there was one pair of identical sequences in 2008, another distinct pair was identified in 2012 which was also detected once in 2015. For Subject 2, there were 3 intact identical sequences identified at the last time point out of a total of 6 intact proviruses. The presence of identical sequences is suggestive of clonal expansion of cells harboring intact proviruses, a recently supported phenomenon which may contribute significantly to the maintenance of the intact reservoir ^14,32–41^. To assess the role of clonal expansion, we next counted each clone only once when it first appeared (Figure 2E-F), assuming proviruses with identical sequences were clones, and found that this led to a greater R-squared value of the fit to the exponential model, suggesting that when the effects of clonal expansion are reduced the resulting dynamics more closely follow an exponential decay.

### Deletion analysis suggests that splice sites play an important role in both negative and positive selection pressures (Figure 3)

We analyzed the selection pressures on defective proviruses by removing all intact sequences, as we reasoned in this setting the potential confounding factor of ongoing replication would be removed. The population of defective proviruses was a mixture of mostly deleted and some hypermutated proviruses by APOBEC (between 5-14% of total)^42–48^. To visually inspect for selective pressures on defective proviruses, we aligned the deletions of defective proviruses to HXB2 for Subject 1 (Figure 3B-C) and Subject 2 (Figure 3D-E) at the first and last time point to examine if any pattern emerged. For visual clarity, we excluded proviruses shorter than 500 bp, proviruses with deletions of less than 100 bp and hypermutated proviruses. We grouped proviruses into categories based on the presence or absence of the donor splice sites 1 (D1) and 4 (D4). D1 and D4 are unique among HIV splice sites for their strong ability to interact with U1 snRNP and splice with a downstream acceptor. The other splice donor and acceptor sites of HIV are all considered weak ^16,49^. We grouped deletions as follows: D1+D4−(red), D1−D4+ (blue), D1−D4− (gold) and D1+D4+ (black). Upon inspection, the D1+D4− (red) appeared to contract over time and the D1−D4+ (blue) appeared to expand over time in Subject 1. Upon studying the red (D1+D4−) deleted proviruses, it was apparent that a large fraction of these proviruses in Subject 1 would be capable of expressing HIV Gag and some could also express HIV Pol providing a mechanism for proviral clearance. In contrast, the proviruses in blue (D1−D4+) had one deletion at the 3’ end. Thus, it became clear that these proviruses would not be able to express HIV proteins efficiently because they lacked the proper AUG for Gag/Pol, and D1, which is utilized in the canonical splice pathway for all of the proteins encoded on the 3’ end of HIV. Thus, the blue proviruses were unlikely to express HIV proteins efficiently and so were less likely to experience negative selection, but rather appeared to experience relative positive selection compared to the red proviruses. Inspection of Subject 2 shows that proviruses capable of making Gag decreased slightly and proviruses capable of strong splicing increased relatively. The less dramatic changes in Subject 2 could be due to weaker immune responses, which in turn would be consistent with his clinical history, characterized by a rapid drop in CD4 T-cell count (down to 0 CD4 T cells six years after diagnosis) and severe opportunistic infections before starting ART.

**Figure 3.**
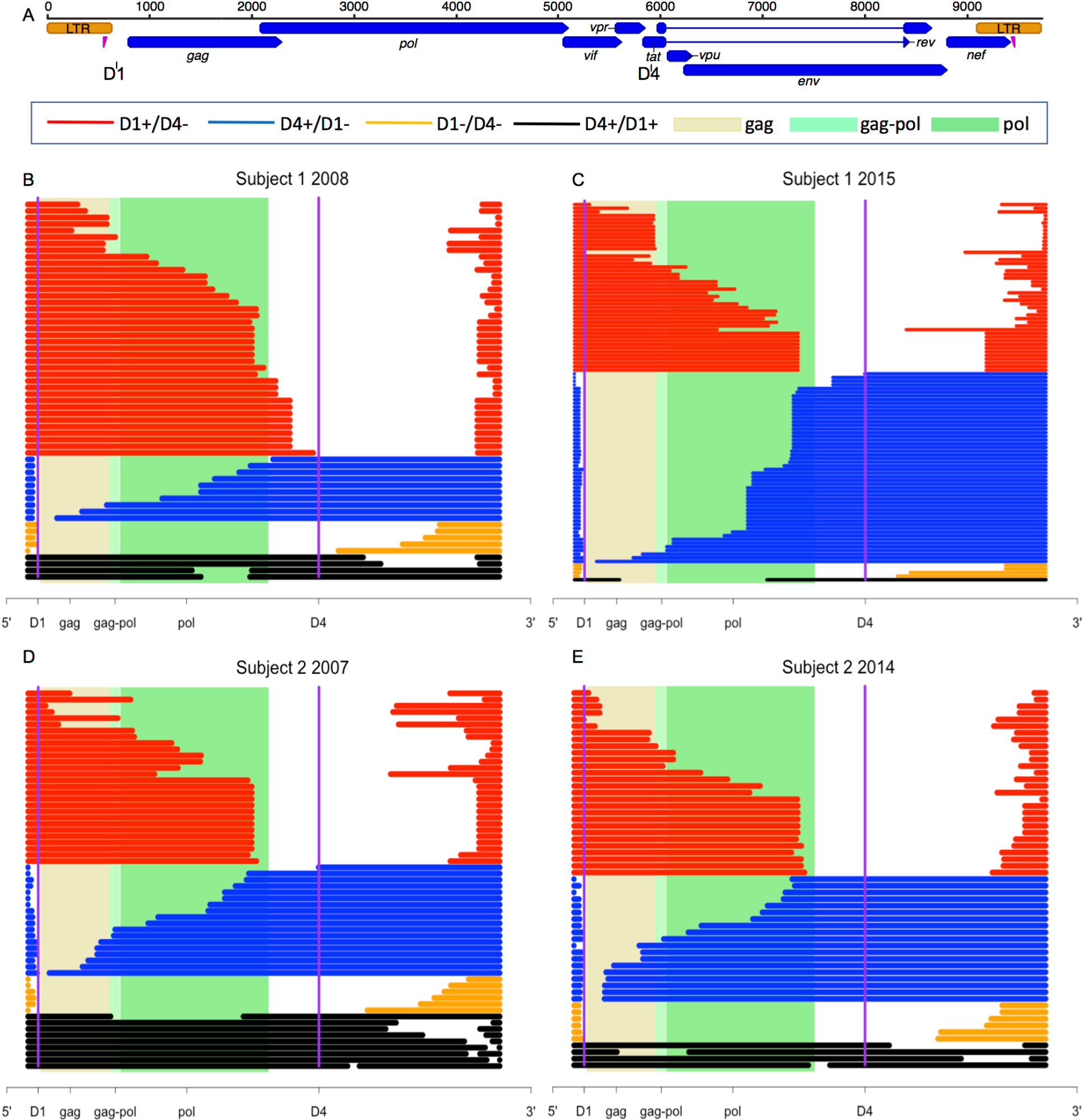
Deletion analysis suggests that splice sites play an important role in both negative and positive selection pressures. A) Structure of HIV genome with genes denoted by the blue bars. Location of strong donor splice sites D1 and D4 are specified. B) Defective proviruses from Subject 1 for the first apheresis sample in 2008 aligned to HXB2. Red proviruses have deleted D4, blue proviruses have deleted D1, both splice sites are deleted in the yellow proviruses, and both splice sites are present in the black proviruses. The shaded beige, light green and dark green regions correspond to the gag, gag-pol and pol regions of HXB2 respectively. C) Defective proviruses from Subject 1 for the last apheresis sample in 2015. Proviruses are graphed on the same scale to demonstrate how the proportion of each type of defective proviruses changed from first to last time point. D) Defective proviruses from Subject 2 for the first apheresis sample in 2007. E) Defective proviruses from Subject 2 for the last apheresis sample in 2014. Proviruses shorter than 500 bp, proviruses with deletions of less than 100 bp and hypermutated proviruses were excluded from this analysis.

### Proviruses that maintain the potential to express HIV proteins experience relative negative selection (Figure 4A)

To probe if expression potential was important for selective pressure we asked if defective proviruses with an intact D1 sequence and at least one intact ORF were selected against. We asked if we could detect a significant decline in D1 splice sites with at least one intact ORF over time. We required the presence of D1 and at least one intact ORF because proteins that are expressed on the 3’ end utilize this sequence to make a spliced message ^16,17^; we found the majority of proviruses meeting these criteria contained an intact Gag ORF (Figure 3). We found that in both Subjects, there was a significant linear decline in D1 splice sites with at least one intact ORF among defective proviruses (Subject 1, p=0.02, R^2^=0.97; Subject 2, p=0.04, R^2^=0.92). Thus, our data suggest that the capability to express proteins even among defectives correlates with relative clearance of proviral DNA.

**Figure 4.**
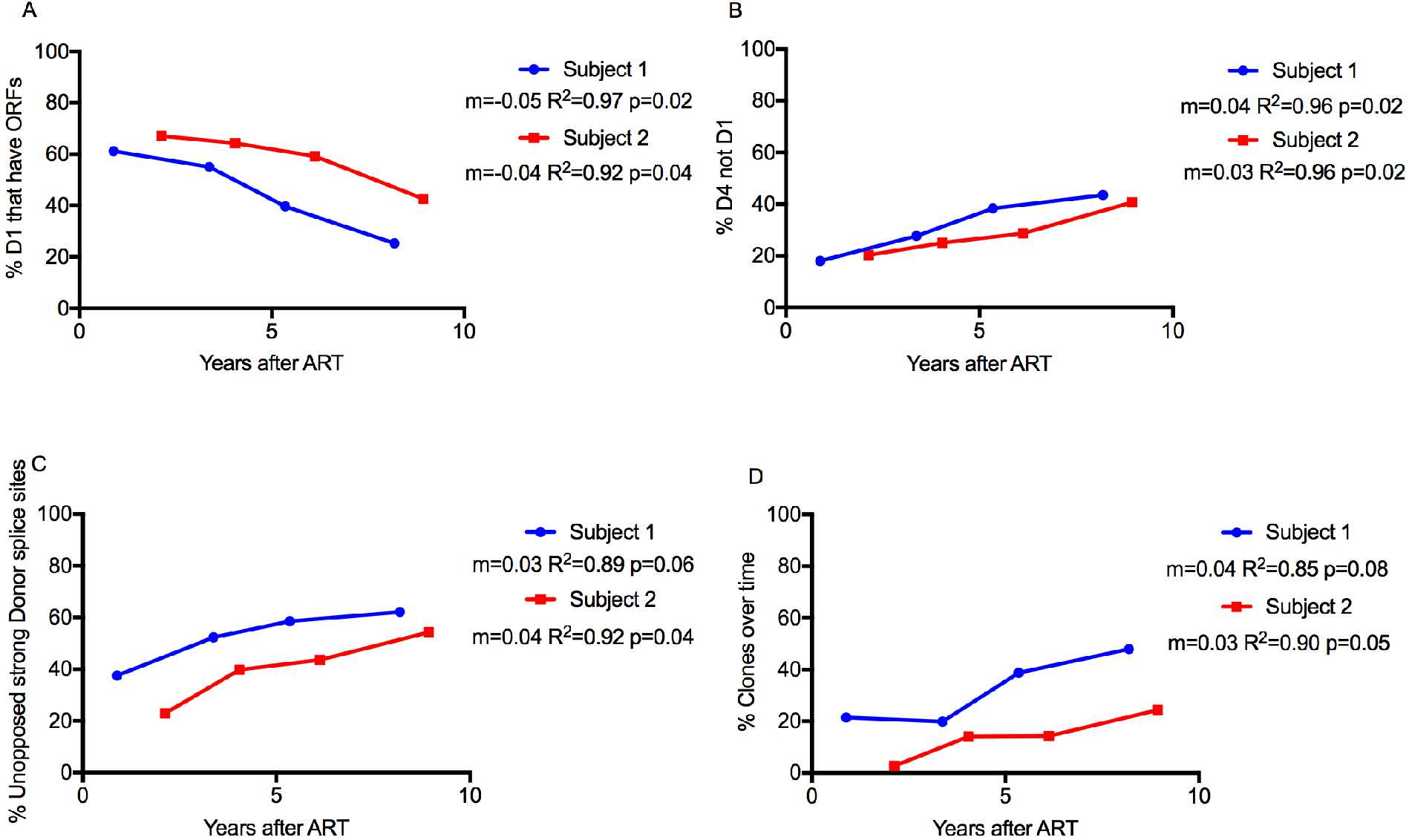
Relative changes in the major 5’ splice sites donors (D1 and D4) and clones in defective proviruses over time. A) Percentage of intact D1 splice site sequence in proviruses with at least one functional ORF at each time point. B) Percentage of intact D4 splice site sequence in defective proviruses lacking 5’ D1 at each time point. C) Percentage of defective proviruses with “unopposed” strong donor splice site, i.e. D1+ without ORFs or D4+ without D1 at each time point. D) Percentage of clones over time in defective proviruses. Statistical values are calculated by Pearson correlation.

### The presence of a strong donor site 4 in the absence of protein expression potential may drive the relative increase in proviruses with deletions in *gag* (Figure 4B)

We next asked if the presence of a strong D4 in the absence of D1 could explain the relative increase of defective proviruses containing ORFs in the 3’ end over time. We reasoned the absence of D1 in a provirus with a large 5’ deletion would prevent expression of high levels of any of the spliced proteins ^16,17^. To test this hypothesis, we plotted the presence of D4 without D1 over time among defective proviruses in both Subjects and found a significant increase (p=0.02, R^2^=0.96 for both), consistent with relative positive selection for proviruses with strong donor splice sites without strong potential for HIV protein expression.

### The presence of a strong D1 or D4 in the absence of protein expression may drive the relative increase in proviruses with deletions overall (Figure 4C)

We then asked if the presence of a strong donor splice site (D1 or D4) in the absence of protein expression might drive the relative increase in deleted proviruses. We defined those proviruses that lack genetic elements to promote protein expression (D4+ without D1 and D1 + without ORFs) as proviruses with an “unopposed” strong donor splice site. In other words, “unopposed” indicates proviruses without genetic elements providing strong potential to express proteins. We found a correlation between the percentage of “unopposed” splice sites and time (p=0.06, R^2^=0.89 for Subject 1 and p=0.04, R^2^=0.92 for Subject 2). In summary, the number of defectives proviruses that contained “unopposed” strong donors increased over time which likely reflects proliferation of cells containing these defective proviruses with strong splice donors.

### Proviral clones are more likely to have strong donor splice sites that lack potential to express HIV proteins than other defective proviruses (Table 1)

We then asked if the presence of a strong donor splice site (D1 or D4) in the absence of protein expression might drive clonal expansion by calculating the percentage of clones that had D1 and no ORF or had D4 but no D1 among the clones as well as among all defective proviruses (Table 1). Comparing columns E and F, at every time point we found more “unopposed” strong donors splice sequence among clones than among defective proviruses.

**Table 1.** Proviral clones are more likely to have strong donor splice sites that lack potential to express HIV proteins.

|  | Apheresis<br>time points | A | B | C | D | E | F |
| --- | --- | --- | --- | --- | --- | --- | --- |
|  |  | D1<sup>a</sup> in<br>clones with<br>no ORFs<sup>b</sup><br>(N) | D4<sup>c</sup> not D1<br>in clones<br>(N) | A+B<br>(N) | Total<br>clones<br>(N) | C/D<br>(%) | Defective<br>proviruses with<br>D1 and no ORFs<br>or D4 but not D1<br>(%) |
| **Subject**<br>**1** | 2008 | 6 | 2 | 8 | 15 | 53 | 38 |
|  | 2010 | 12 | 6 | 18 | 25 | 72 | 52 |
|  | 2012 | 9 | 22 | 31 | 38 | 82 | 59 |
|  | 2015 | 9 | 40 | 49 | 59 | 83 | 62 |
| **Subject**<br>**2** | 2007 | 0 | 1 | 1 | 2 | 50 | 23 |
|  | 2009 | 2 | 4 | 6 | 12 | 50 | 40 |
|  | 2011 | 1 | 12 | 13 | 13 | 100 | 44 |
|  | 2014 | 1 | 11 | 12 | 19 | 63 | 54 |

<sup>a</sup>D1, Donor splice site 1
<sup>b</sup>ORFs, Open Reading Frames
<sup>c</sup>D4, Donor splice site 4

### Defective clones increase relatively over time (Figure 4D)

We plotted the frequency of all clones over time as a proportion of the reservoir. Subject 1 had more clonal proviruses than Subject 2. In both Subjects proviral clones expanded linearly and the rate of clonal expansion was similar (the slope describing clonal expansion for Subject 1 was 0.04, p=0.08 and the slope for Subject 2 was 0.03, p=0.05). The slope was statistically greater than 0 for Subject 2 and trended to be positive in Subject 1, with clones reaching nearly 60% of the reservoir at the last time point. Clones steadily increased relative to other defective proviruses over time in both Subjects, consistent with several recent studies ^35–38,50^.

### Clonal expansion may occur before ART initiation (Figure 5)

In Figure 4 (A-D) we analyzed the relative contribution of positive and negative selection on the proviral landscape over time. We based our choice to use relative changes on previous work showing that HIV DNA is constant over time on ART ^7^ and our own data showing that HIV DNA did not change more than 2-fold over the study period. We also decided to analyze the absolute change in the number of defective proviruses over time (Figure 5 A-D). We observed a decline in the number of defective proviruses containing D1 and functional ORFs for Subject 2 (p=0.05, R^2^=0.91) and a trend toward decline for Subject 1 (p=0.20, R^2^=0.65) (Figure 5A). This suggests the clearance of proviruses with the potential to express Gag/Pol. We did not observe a clear pattern of positive selection for the absolute level of defective proviruses with strong donor splice site and poor potential for protein expression (Figure 5B-D). One explanation for lack of significance is that the error introduced by our PCR assay is obscuring a positive slope. Given the high potential for PCR error for DNA measures due to sequence variation and that DNA measures never differed more than 2-fold from any time point, we refrain from deep interpretation of these data. One explanation deserving further analysis is that clonal expansion happened in large part before ART initiation and is revealed by clearance of other proviruses expressing proteins when T-cell turnover is higher ^51^.

**Figure 5.**
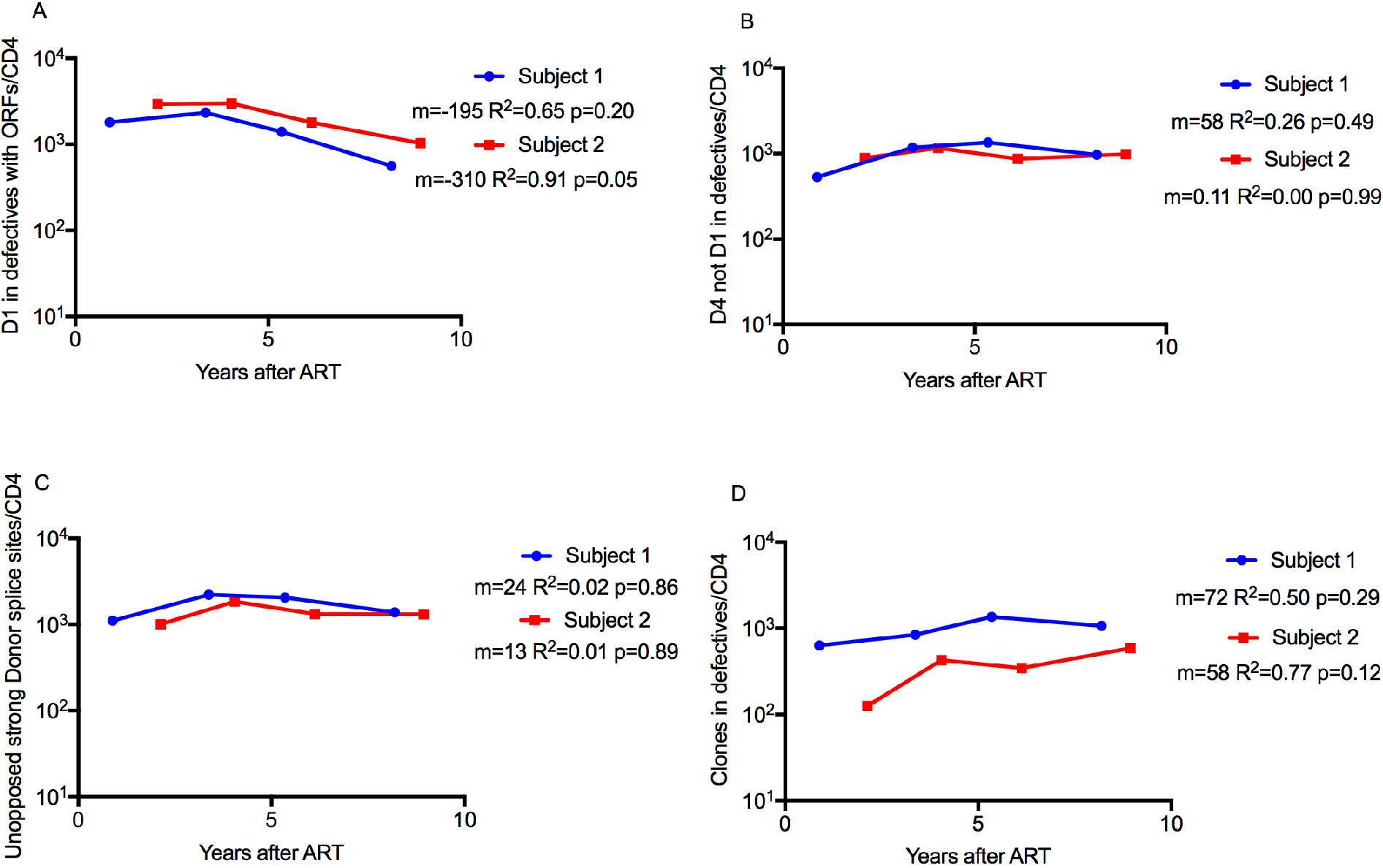
Absolute changes in the major 5’ splice sites donors (D1 and D4) and clones in defective proviruses over time. A) Absolute number of intact D1 splice site sequences in proviruses with at least one functional ORF at each time point. B) Absolute number of intact D4 splice sequences in defective proviruses lacking 5’ D1 at each time point. C) Absolute number of defective proviruses with “unopposed” strong donor splice site, i.e. D1+ without ORFs or D4+ without D1 at each time point. D) Absolute number of clones over time in defective proviruses. Statistical values are calculated by Pearson correlation. Absolute numbers are normalized per CD4 T-cell count.

## Discussion

Our longitudinal study of proviral sequences reveals that two opposing forces simultaneously exert negative and relative positive selection pressures on proviral DNA in HIV-infected individuals on ART. Specifically, we found that proviruses with the genetic potential to express HIV proteins declined over time while proviruses with strong donor splice sites but limited potential to express HIV proteins increased relatively over time. Consistent with recent papers ^14,34,36,38,41^, we also found evidence of clonal expansion of intact proviruses though this expansion may have in large part occurred before ART initiation. We speculate that positive selection of these proviruses can be driven by the unique position of strong HIV donor splice sites in an intron adjacent to the exon of an oncogene. Our work has several implications for HIV eradication: first, it provides a mechanism for how intact proviruses can decline over time while proviral DNA levels remain unchanged; second, it gives new insights on forces that might drive clonal expansion; third, it provocatively suggests that the HIV reservoir is likely less resistant to reactivation than generally thought. This has important implications for HIV cure, as it argues against the current dogma of “shock and kill” strategies that the major hurdle to HIV eradication is represented by the “invisibility” of the reservoir.

We found that intact proviruses contracted more rapidly than defective proviruses. This finding suggests that intact proviruses experience stronger negative selection. The strong negative pressure against intact proviruses in turn suggests that the majority of the replication-competent reservoir is expressed over time, despite the small fraction that is detectably expressed at any one moment ^39,52–58^. Selective negative pressures could be due to immunity or viral cytotoxicity. In addition, we found that defective proviruses with the potential to express HIV proteins declined more rapidly than other defectives. The fact that most of those cleared proviruses expressed Gag, a relatively nontoxic viral protein, suggests that immune pressure likely plays a role in proviral clearance even during ART. It is generally thought that immune pressure during ART is minimal due to the dramatic drop in the total antigen load ^59–61^. To the best of our knowledge, our study is the first to show that immune pressure can play a role in shaping the reservoir even during suppressive ART in humans. In rhesus macaques there is evidence that CD8 T-cell depletion after ART suppression results in rebound viremia which is consistent with our interpretation ^62^.

Our analysis of defective proviruses provides further evidence that expression of HIV proteins is an important driver for HIV clearance. Specifically, proviruses with a functional gag ORF appear to be cleared over time (Figure 3, 4A and 5A). Studies from Anderson, Maldarelli and colleagues showed by qPCR that HIV gag DNA generally declines over time in subjects on long-term ART (Anderson CROI 2017) consistent with our findings. Notably, Imamichi (IAS 2017) was able to detect HIV Gag from T cells that were cloned from an ART-treated subject. The provirus that was cloned from these T cells contained a 2.4kb deletion that eliminated *tat/rev* and *env*. Thus, it is reasonable to conclude that even defective proviruses that express HIV Gag could undergo negative selective pressure, as seen in our study.

Our data suggest indirectly that integration of certain forms of defective proviruses can trigger positive selection pressures. We speculate that positive selection would be favored when a defective provirus inserts into an intron next to an oncogene exon, and LTR transcription occurs with minimal HIV protein expression. Splicing between a strong HIV donor site and an oncogene acceptor site could result in higher expression of an oncogene as recently described ^63^. Our work advances the findings of Cesana by showing that unopposed strong donor splice sequences correlate with proviral expansion. It also provides a potential mechanism behind other studies showing positive selection for HIV integration near an oncogene ^50,64–66^. In fact, we speculate that providing a strong donor splice sequence might be a quantitatively significant mechanism for clonal expansion (Table 1). Moreover, our work also reveals that viral protein expression may be the substrate for negative selection pressure and may counterbalance the positive selection potential of unopposed strong donor splicing. For example, a provirus that lacks the major donor site D1 should not be able to make HIV proteins except for Gag and Pol as all other HIV proteins are made from spliced HIV RNA which requires D1. Moreover, a provirus that lacks D1, but has D4 is typically deleted in the gag-pol region and thus is unlikely to express any HIV proteins, but it should retain the ability to splice to the adjacent exons of a nearby oncogene. This is supported by work of several groups showing that U1 snRNP can suppress cryptic cleavage and polyadenylation in introns ^67,68^. In this setting, there would be enhanced expression of the oncogene and enhanced cellular proliferation, but there would be little HIV protein expression to exert negative selection. Notably, if HIV inserts within an intron of an oncogene and HIV protein expression is robust, this could lead to competing positive and negative selection pressures within the same cell. Positive selection would occur if a strong D1 or D4 splice sequence can splice to the acceptor of the oncogene leading to forced expression of the oncogene, but on the other hand negative selection pressure may occur if the provirus is intact or capable of expressing HIV proteins that could elicit an immune response. Thus, our study of defective proviruses reveals a possible mechanism for clonal expansion of both defective and intact proviruses. One limitation of our approach is that we cannot precisely quantify the extent of negative and positive selection because they are opposing forces. In addition, we cannot define whether the expansion of clones happened before, after ART or throughout infection. In one scenario, clones might form predominantly before starting ART, then emerge after clearance of the proviruses capable of expressing proteins. This is consistent with previous work showing turnover of T cells is many fold higher before ART is initiated ^51^. Regardless, the enrichment of clones with identical sequence indicates that some proviral expansion occurs through cell division. Our contribution is to provide evidence that a substantial driving force may be due to unique positioning of splicing sites.

Clonal expansion is a newly identified force driving HIV persistence. In our study, we observed that 50% of intact proviruses in Subject 2 were identical at the last time point. This suggests that cell division contributed to the increase in intact proviruses, though we cannot rule out the contribution of a limited amount of viral replication without sequencing the integration sites. Evidence supporting a role for clonal expansion in proviral persistence has been mounting, including that infected effector memory (EM) T cells increase as KI67 positive cells become more prominent ^41^ and that monotypic virus increases over time on ART, as shown in our paper and others ^32,35–38,41,50,63^. Several studies have also shown identical sequences over time - consistent with clonal expansion of the “true” reservoir ^32–38,41,50,63 36,64^. In one study the identical intact sequences were more prominent in EM cells ^10^ consistent with work showing the persistence over many years of a mutant HIV clone in EM cells ^69^. The accumulation of intact clonal sequences (Figure 2) in Subject 2 adds to the evidence that clonal expansion plays a role in reservoir persistence and suggests that its contribution is substantial.

Taken together, our data suggest that intact HIV proviruses are under stronger negative selective pressure for clearance than defective proviruses. This suggests indirectly that the majority of the reservoir is expressed over several years, implying that lack of HIV expression may not be the main hurdle to reservoir clearance, while clonal expansion may represent a new target for HIV eradication.

## Methods

### Apheresis

Subjects underwent apheresis at the University of Pennsylvania according to protocols, #704904, approved by the Institutional Review Board (IRB). The early time point samples from Subject 1 and 2 were provided by Dr Stephen Migueles (National Institute of Health) who follows his institutional protocol with IRB approval.

### DNA Isolation and Quantification of HIV DNA

DNA was isolated from PBMCs using the Gentra Puregene Cell Kit (Qiagen). HIV DNA was quantified by total HIV against the LTR or gag regions and integrated HIV DNA as previously described with minor modifications ^70,71^. qPCR reactions were cycled on a 7500 FAST real-time instrument (ThermoFisher). First step PCR reactions were cycled using the Nexus Master Cycler (Eppendorf).

### Provirus Amplification and Sequencing

A two-step nested PCR approach was used to reduce non-specific amplification from genomic targets. Primer sets used in both reactions were located within the LTRs and were staggered appropriately to avoid localized LTR amplification while simultaneously capturing nearly the full-length of HIV proviruses. For the first PCR reaction we used the following primers: 5’-CCTCAATAAAGCTTGCCTTGAGTGC-3’ and 5’-CCTAGTTAGCCAGAGAGCTCCCAG-3’, HXB2 position 523-547 and 9568-9591 respectively. For the second PCR reaction we used the following primers: 5’-AAGTAGTGTGTGCCCGTCTGTTGTGTGAC-3’ and 5’-GGAAAGTCCCCAGCGGAAAGTCCCTTGTAG-3’, HXB2 position 551-579 and 9429-9458 respectively. We used a long-range and high-fidelity polymerase enzyme for both reactions (Platinum SuperFi PCR Master Mix, ThermoFisher). In the first PCR reaction, PBMCs DNA was diluted so that PCR amplification resulted in ≤ 30% of wells being positive for HIV DNA. Nested PCR reactions were visualized by gel electrophoresis, and the fraction of reactions containing ≥ 2 bands were excluded from our analysis as these were often found to contain multiple proviruses. PCR amplicons were purified using the DNA Clean & Concentrator kit (Zymogen) and DNA concentration was measured using the Quant-iT dsDNA Broad Range Assay Kit (ThermoFisher). Amplicons were prepared using the Nextera library preparation kit (Illumina) and sequenced on a MiniSeq System using a Mid-output flow cell (Illumina).

### Sequence Assembly

Paired reads were trimmed in the program Geneious using the BBDuk plugin, discarding reads from the adaptor, and then merged using Geneious. Again, the reads were trimmed of those with a quality rating under 30, and those under 115 base pairs in length. The reads were then mapped using BBMap to the HXB2 reference sequence, and the reads which aligned to the HIV sequence were extracted. Provirus contigs were made through *de novo* assembly of the extracted reads. Contigs generated by Spades, Tadpole, and Trinity ^72^ *de novo* assemblers were compared. Accuracy of each assembler was evaluated by: 1) its ability to produce a contig which matched the length of the region supported by reads when mapped to an HIV reference 2) reads supporting the generated contig. We selected Spades as our default *de novo* assembler based on these criteria.

Reads were *de novo* assembled using Spades with default settings. When reads gave rise to multiple non-overlapping contigs, the contigs were concatenated into one sequence. The final contig was then mapped back to HXB2, and annotated with motifs, including splice donor and acceptor sites, and open reading frames (ORFs) as described in supplementary methods. Finally, in order to determine whether two proviruses had been sequenced together into one contig (double proviruses), the extracted reads from before were aligned to the assembled contig.

### Removal of Double Proviruses

Double proviruses were identified according to criteria listed in the supplementary methods and discarded from analysis.

### Nomenclature

Intact proviruses were defined as those determined to code for nearly psi packaging sites with at least 3 stem loops (SL2 has to be intact because it contains D1 and a cryptic donor site that works when the Major Donor Site 1 is mutated ^17^), and 9 complete ORFs for all HIV genes. We allowed for truncated Nef and Tat genes as commonly identified in infectious strains of HIV. Nef was allowed to be truncated up to the extent seen in NL4-3. We required the presence of Major Donor Site 1 or a GT dinucleotide cryptic donor site located four nucleotides downstream ^17^(only found in three proviruses) and presence of Major Donor Site 4. We also required the presence of splice Acceptor Site A3 (A4), A5, A7, either A4a or A4b or A4c as well as an intact Rev-responsive element (RRE) sequence. Our rationale for these criteria is described ^17^. We also accepted the sequence “GGTAAGT” as well as the canonical donor 1 splice sequence “GGTGAGT” for the D1 sequence as these sequences binds U1 snRNP equally well (Personal communications C. Martin Stoltzfus). Notably this D1 variant sequence was found in a proviral sequence with no functional ORFs that was present at increasing frequency over time, consistent with clonal expansion.

### Intact Provirus Decay Analysis

Based on intact criteria, the number of intact proviruses per million CD4 T cells was calculated for each patient at the four apheresis time points. To estimate decay parameters, a statistical analysis was performed using a random-effects regression model assuming first order decay kinetics. Setting time t=0 to be the date when ART was initiated, we are able to estimate the number if intact proviruses at the beginning of treatment, decay rate, as well as their half-life.

### Deletion Analysis

Provirus consensus sequences were aligned by MAFFT ^73^ using the iterative E-INS-i method with a gap penalty opening penalty of 1.8. This facilitates proper alignment of proviruses of different lengths, which is common among proviruses with deletions. Aligned sequences were then exported to an R software environment using the Seqinr Biological Sequence Retrieval and Analysis package. Once in the R software environment a program removed base pairs within patient proviruses which were insertions relative to the HXB2 HIV sequence. This allowed the alignment of all proviruses to be standardized in length and base pair indices to HXB2. Then, deletions with length more than 100 base pairs were recorded, and a graph was made in ‘R’ which showed each sequence plotted against the base pair numbers of HXB2, with deletions removed (Figure 3).

### Identification of Hypermutant sequences

To identify hypermutant HIV sequences, all proviruses for each patient were aligned using MAFFT with the E-INSi algorithm and a 1.8 gap penalty, and an intact HIV sequence was selected as the reference. The aligned proviruses were checked against the reference for hypermutants using the LANL Hypermut 2 program. The provirus with the lowest chance of being a hypermutant as determined by hypermut was selected as the reference, and once again Hypermut 2 was run on the alignment. Proviruses determined to be hypermutant with p<0.05 were counted as hypermutant ones.

### Phylogenies and Identification of Potential Clones

Intact proviral sequences were aligned using MAFFT ^73^ with the G-INSi algorithm with a 1.8 gap penalty. A maximum likelihood tree was constructed using PHYML with the general time reversible substitution model, using both SPR and NNI optimization methods for topology, 4 substitution rate categories, and an estimated transition/transversion ratio, proportion of invariable sites, and gamma distribution parameter ^74^.

As described above, all proviruses (both intact and defective) were aligned in MAFFT using the E-INSi algorithm to find potential clones, defined as proviruses with the same sequence and similar length. This was first done by creating a phylogeny of all proviruses as described above, except those with inversions and large insertions. Proviruses that clustered closely in the phylogeny were then individually aligned with each other and manually checked for identical sequences (with sequence differences highlighted by Geneious). Clones were checked for a second time, this time once again aligning all proviruses with MAFFT and then among proviruses of a similar length manually identifying proviruses of identical sequence.

### Statistical Analysis and Graphing

Statistical processes were performed using ‘R’ and Microsoft Excel, and MATLAB’s Prism software was used for graphing.

## SUPPLEMENTAL METHODS

### Criteria for Double Proviruses

#### ANY OF

1. One or more point mutations varying between two alternatives with the following exceptions:
  a. Two or less point mutations which vary between alternatives within 40 bp of the start/end of the first/end read of the entire sequence
  b. A single point mutation varying between two alternatives where one of them is present less than 40% of the time, or is present in near exactly a whole multiple of two times less often than the alternative
  c. Insertions/deletions of A nucleotides at the beginning/end of chains of at least 5 consecutive A nucleotides
2. A region at least 500 basepairs (bp) long where coverage suddenly drops at least 45%
3. A combination of exception 1b or 1a and at least 1000 bp where coverage is under 50%
4. For deletions spanned by some proviruses but not others, a provirus is considered as double if a deletion is accompanied by a point mutation, or if the less common version occurs at least 30% of the time. However, if the sequence spanning the deletion in the sequences that do not have the deletion is identical to sequences elsewhere in the provirus, or if the deletion is only spanned or deleted by proviruses where the read just barely ends just into or soon after the deletion, it is considered an assembly error and only one provirus

### Motifs and ORFs Identification

ORFs were determined through Geneious, defined as at least 50 nucleotides following ‘ATG’ or 3000 following ‘TTT’ (for the Pol ORF), and ending with a stop codon. When mapped to HXB2, ORFs in the proviruses were matched to specific HIV ORFs if they were within 20 nucleotides. To identify Tat and Rev, parts 1 and 2 of Tat and Rev were mapped to the provirus based on 65% homology with the HXB2 Tat and Rev 1 and 2. These Tat 1/2 and Rev 1/2 homologous sequences of the provirus were then extracted and the 1 and 2 parts concatenated and translated. The homologous Tat and Rev were determined intact if the sequences had no early stop codons and a stop codon in the correct location.

To identify splice and packaging sites, motifs were annotated using Geneious. They were identified as specified in the table below. Note that one of the D1 sequences also occurred later in the HIV genome, and these extraneous annotations were removed.

**Supplemental Table 1.**
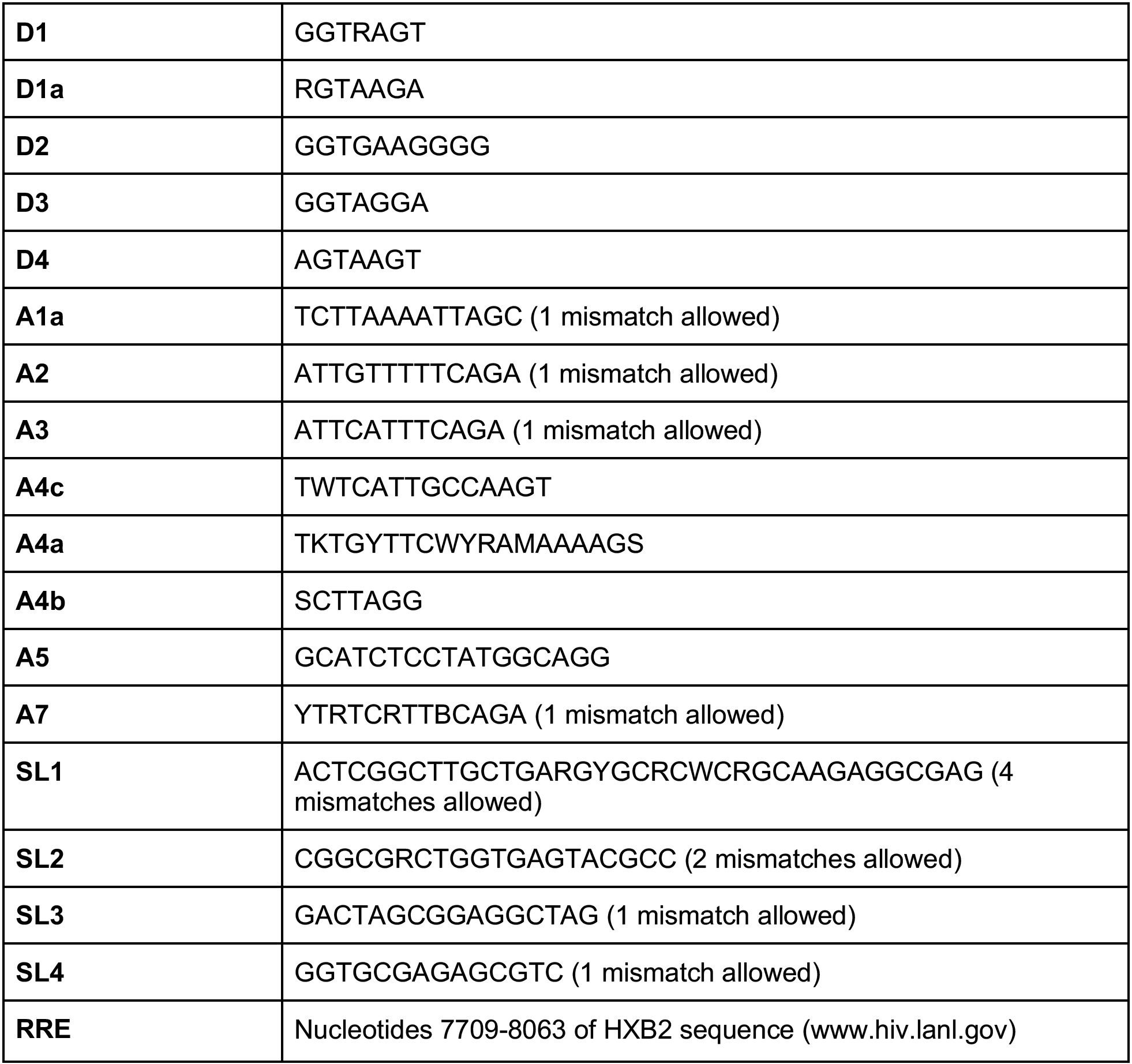
List of the sequences of splice and packaging sites used to identify intact proviruses.

**Supplemental Figure 1.**
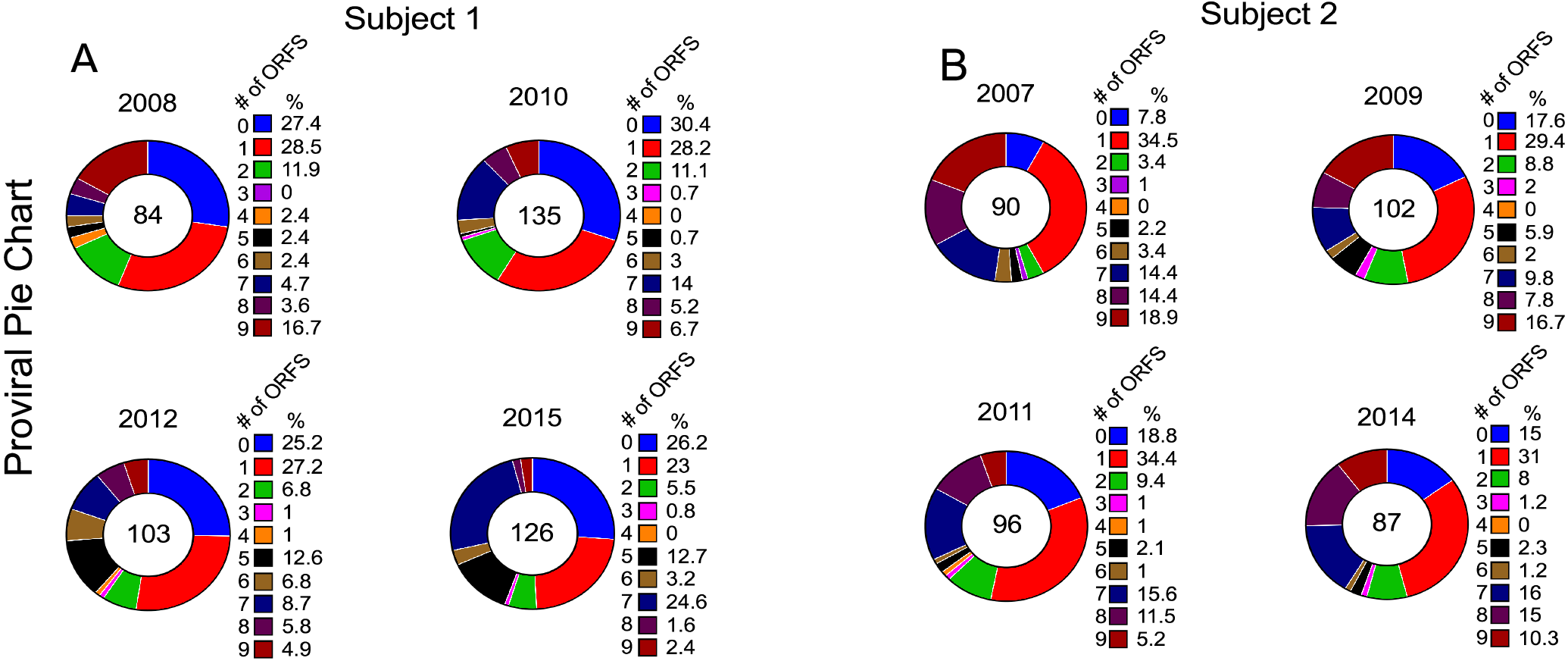
Number of Intact Open Reading Frames over time in Subject 1 (A) and Subject 2 (B). The total number of proviruses sequenced at each time point is reported in the center of the pie chart. For each time point, we also report the number of proviruses containing from 0 to 9 ORFs as percentage over time.

## Acknowledgments and funding

We would like to thank our two Subjects for their participation in the study. We would also like to thank Ryan Zurakowski, Anastasios Vourekas, Martin Stoltzfus and Giuseppe Nunnari for their intellectual contribution.

This work was supported by the National Institute of Allergy and Infectious Diseases of the National Institutes of Health under award numbers R01AI12001, R21AI116216, UM1AI126617, R33AI104280 with co-funding support from the National Institute on Drug Abuse, the National Institute of Mental Health, and the National Institute of Neurological Disorders and Stroke. The content is solely the responsibility of the authors and does not necessarily represent the official views of the National Institutes of Health.

## Author contributions

M.R.P., D.J.V., S.W., M.P.B. and U.O. designed the experiments. M.R.P and D.J.V. performed HIV DNA quantification, proviral amplification and sequencing experiments. S.W., M.P.B., L.C., B.S., and W.H. performed data analysis. U.O. and M.R.P. wrote the manuscript. M.R.P., D.J.V., S.W., and M.P.B contributed equally to the work.

